# Autophagy mediates cancer cell resistance to doxorubicin induced by the Programmed Death 1/Programmed Death Ligand 1 immune checkpoint axis

**DOI:** 10.1101/2023.01.17.524124

**Authors:** Lori Minassian, Daniel Sanwalka, Jean-François Paré, Shannyn Macdonald-Goodfellow, R. Liam Sutherland, Abdi Ghaffari, Chelsea L. Margerum, Madhuri Koti, Andrew W.B. Craig, Tiziana Cotechini, D. Robert Siemens, Edmond Chan, Charles H. Graham

## Abstract

**Background:** While the Programmed Death 1/Programmed Death Ligand 1 (PD-1/PD-L1) immune checkpoint is an important mechanism of immune evasion in cancer, recent studies have shown that it can also lead to resistance to chemotherapy in cancer cells via reverse signaling. Here we describe a novel mechanism by which autophagy mediates cancer cell drug resistance induced by PD-1/PD-L1 signaling.

**Methods:** Human and mouse breast cancer cells were treated with recombinant PD-1 (rPD-1) to stimulate PD-1/PD-L1 signaling. Activation of autophagy was assessed by immunoblot analysis of microtubule-associated protein 1A/1B-light chain 3 (LC3)-II and Beclin 1 protein levels, two important markers of autophagy. Moreover, autophagosome formation was assessed in human breast cancer cells using green fluorescence protein (GFP)-tagged LC3. Cells were either treated with Beclin 1 or Atg7 shRNA to assess the role of autophagy on resistance to doxorubicin mediated by PD-1/PD-L1 signalling. We then investigated signaling mechanisms upstream of PD-1/PD-L1 induced autophagy by assessing phosphorylation of extracellular signal-related kinase (ERK).

**Results:** Treatment of cells with rPD-1 resulted in a time-dependent increase in LC3-II as well as Beclin 1, and an increase in autophagosome formation. Knockdown of Beclin 1 or Atg7 prevented drug resistance induced by PD-1/PD-L1 signaling. Exposure of breast cancer cells to rPD-1 resulted in increased ERK phosphorylation and inhibition of ERK activation abolished autophagy induced by PD-1/PD-L1 signaling.

**Conclusions:** These studies provide a rationale for the use of PD-1/PD-L1 immune checkpoint blockers and autophagy inhibitors as potential chemosensitizers in cancer therapy.

## INTRODUCTION

An important immune checkpoint in cancer involves the upregulation and binding of Programmed Death Ligand 1 (PD-L1) on cancer cells to Programmed Death 1 (PD-1) on cytotoxic T cells. This ligand-receptor interaction leads to T cell anergy, exhaustion, apoptosis, and decreased cytokine production and proliferation, ultimately resulting in decreased cytotoxic activity against tumor cells (1). Thus, much effort has gone into developing PD-1/PD-L1 immune checkpoint blockers, some of which have demonstrated unprecedented clinical efficacy as first-line or second-line therapy of various cancers including melanoma, non-small cell lung cancer, renal cell carcinoma, urothelial carcinoma, and colorectal cancer, among others (2–5).

Because the PD-1/PD-L1 axis is an important mechanism of immune escape, most studies have focused on evaluating how PD-1-mediated signaling affects T cell properties. However, recent reports have demonstrated that PD-1/PD-L1 signaling is bi-directional (6,7) and that engagement of PD-1 with PD-L1 can result in acquisition of drug resistance in tumor cells (8). We recently discovered that, following exposure to recombinant PD-1, breast and prostate cancer cells that express PD-L1 acquired resistance to different classes of conventional anti-cancer drugs, *i.e.* doxorubicin and docetaxel. Moreover, acquisition of this drug resistance phenotype was prevented when the PD-1/PD-L1 interaction was inhibited by incubating cells with anti-PD-1 antibody, anti-PD-L1 antibody or PD-L1 targeting siRNA (8). That study, however, did not fully characterize the mechanism of drug resistance.

The autophagy intracellular degradation pathway plays a number of well-established roles in sustaining the high metabolic levels required for the abnormal growth of cancer cells (9). In addition, autophagy has also been proposed to contribute to drug resistance mechanisms following therapy with various agents such as 5-flurouracil, cisplatin, gemcitabine, anthracyclines, and tamoxifen (9–15). In several contexts, inhibition of autophagy enhances the efficacy of chemotherapy (12–14). Thus, in the present study, we investigated the contribution of autophagy to breast cancer cell drug resistance induced by PD-1/PD-L1 signaling. We report here that engagement of PD-1 with PD-L1 on tumor cells mediates drug resistance via induction of autophagy.

## MATERIALS AND METHODS

### Cell Lines

Human MDA-MB-231 breast cancer cells and mouse mammary carcinoma 4T1 cells were purchased from the American Type Culture Collection (Manassas, VA, USA) and were cultured in RPMI 1640 medium (Wisent 350-000-CL; Wisent Bioproducts, Montreal, QC, Canada) supplemented with 5% or 10% fetal bovine serum (FBS; Sigma #F6178; Sigma-Aldrich Canada, Oakville, ON, Canada), respectively.

### Flow Cytometry to assess PD-L1 expression

Flow cytometry was performed according to a standard protocol from Abcam (Toronto, ON, Canada). Cells were stained for one hour using the following antibodies: Allophycocyanin (APC) anti-human CD274 (5 μL/test; BioLegend #329707; BioLegend, San Diego, CA, USA), APC anti-mouse CD274 (5 μL/test; BioLegend #124311), and matched isotype controls APC mouse IgG2b k isotype control (5 μL/test; BioLegend #400322), APC Rat IgG2b, k Isotype Control (5 μL/test; BioLegend #400611) and then fixed with 2% paraformaldehyde. Staining was assessed on the MACS Quant Analyzer (Miltenyi Biotec, Bergisch Gladbach, Germany) and analysis was done with FlowJo™ Version 10 (BD, Franklin Lakes, NJ, USA).

### Exposure to rPD-1

Tumor cells were seeded in six-well plates at a density of 2 x 10^5^ cells/well in RPMI 1640 medium supplemented with 2% FBS. Twenty-four hours later, cells were treated with 1 μg/mL of either human rPD-1 (hrPD-1-Fc; R&D Systems #1086-PD-050; Minneapolis, MN, USA) or mouse rPD-1-Fc (R&D Systems #1021-PD-100) for 0, 10, 30, 60 and 180 minutes. In some experiments where indicated, 30 minutes prior to addition of rPD-1-Fc (dosage according to Cell Signaling Technologies (CST) protocol; Danvers, MA, USA) cells were incubated with anti-PD-L1 blocking antibody (Ab; 4 μg/mL; BioLegend #29702; dosage determined from previous work (8)), or the mitogen-activated protein kinase kinase (MEK) inhibitor U0126 (10 μM; CST #9903).

### Immunoblotting

Proteins from cell extracts were separated by sodium dodecyl sulphate-polyacrylamide gel electrophoresis (SDS-PAGE) on 4-20% gradient gels (Bio-Rad #4561095; Hercules, CA, USA) and transferred to polyvinylidene difluoride (PVDF) membranes. Membranes were blocked in 5% skim milk or BSA in 1x Tris-buffered saline/0.1% Tween-20 (TBST) for one hour at room temperature prior to overnight incubations at 4°C with the following primary antibodies: rabbit anti-LC3B (Novus NB11-2220, 1:1000; Littleton, CO, USA), rabbit anti-Beclin1 (Novus NB500-249, 1:1000), mouse anti-pERK (Santa Cruz sc7383; Dallas, TX, USA; 1:500), rabbit-anti-ERK1 (Santa Cruz sc-94, 1:1000), rabbit anti-α-actinin (Santa Cruz sc-15335, 1:1000), or mouse anti-β-actin (Sigma Aldrich A5441, 1:10000). Secondary antibody incubations were performed for one hour at room temperature with the following antibodies: goat-anti-rabbit IgG HRP (Abcam ab97051) and goat-anti-mouse (Bio-Rad #1706516). Protein bands were visualized using chemiluminescence substrate (PerkinElmer #NEL104001EA; Waltham, MA, USA) and the Azure Biosystems c600 Gel Imaging System (Dublin, CA, USA. Densitometric analysis was performed using ImageJ version 1.51k.

### Autophagosome Formation and Imaging

MDA-MB-231 cells were seeded in eight-well chambered polymer coverslips (ibidi #80826) at a density of 4.5 x 10^3^ cells/well. After 24 hours, cells were transfected with 0.24μg EGFP-LC3 plasmid DNA (Addgene #11546; Watertown, MA, USA) using the FUGENE6 transfection reagent (Promega, Madison, WI, USA) in Opti-MEM medium (ThermoFisher). Eighteen hours after transfection, cells were treated with rPD-1 in the absence or presence of anti-PD-L1 antibody or U0126 as described above. After treatment, cells were fixed with 100% cold methanol and imaged with a spinning disk confocal microscope (Quorum WaveFX; Quorum Technologies, Puslinch, ON) at 40x magnification. Ten cells per condition were selected randomly by an observer blinded to the treatment conditions. Autophagosomes were quantified using Image-Pro Plus and ImageJ version 1.51k. Data were pooled from at least three independent experiments.

### Knockdown of Beclin 1 and Atg7 expression

Knockdown of Beclin 1 (using pLKO.1 for BECN1 clone TRCN0000087290) or Atg7 (using pLKO.1 for Atg7 clone TRCN0000092163) in mouse 4T1 cells was achieved following seven days of selection in puromycin as previously described (16,17). pLKO scrambled shRNA (Addgene (#1864)) was used as control.

### Clonogenic assays

Clonogenic assays were performed to determine the effect of autophagy on PD-1-induced drug resistance. 4T1 cells expressing shRNA to knock down Beclin 1 or Atg7 expression or control 4T1 cells expressing scrambled shRNA were treated with mouse rPD-1-Fc (1 μg/mL) for 30 minutes and subsequently incubated in medium containing doxorubicin (0.5 μM; Sigma #D1515) for one hour. The concentration of doxorubicin chosen was based on preliminary dose-response survival data. Cells were then trypsinized, plated at 300 cells/well, and colony numbers were counted 10 days later. The plating efficiency (PE) of all treatment groups was calculated by dividing the number of colonies counted by 300 (number of cells plated). Surviving fraction (SF) was calculated by dividing the PE of cells in all treatment groups by the mean PE of untreated (shRNA alone) control cells. Results from two independent experiments conducted in replicates of three wells per condition in each experiment were pooled for a total of six replicates.

### Statistical Analysis

Statistical analysis was performed using GraphPad Prism software (versions 7.02 and 9.4.0; GraphPad Software, San Diego, CA, USA). Error bars represent standard error of the mean. Two-tailed unpaired Student’s t-test was used to compare means of two groups. One-way analysis of variance (ANOVA) followed by Tukey’s post-hoc test was used when comparing means of three or more groups. Two-way ANOVA was used for experiments involving two independent variables. Data were considered statistically significant when p<0.05.

## RESULTS

### Human and mouse cancer cells exposed to rPD-1 express increased levels of the autophagy markers, LC3-II and Beclin 1 in vitro

MDA-MB-231 and 4T1 cells were confirmed to express PD-L1 (Figure 1A). Conversion of LC3-I to LC3-II and global levels of Beclin 1 are established measures of autophagy. MDA-MB-231 cells treated with 1 μg/mL rPD-1 exhibited a time-dependent increase in lipidated LC3-II (Figure 1B), which led to a statistically significant (p<0.05) LC3 activation ratio (LC3-II/actin) at the 30-minute timepoint. Thus, subsequent incubations involving the use of rPD-1 to stimulate autophagy involved a 30-minute incubation time. MDA-MB-231 cells treated with rPD-1 also showed a time-dependent increase in Beclin 1 levels (p<0.001), peaking at 60 minutes following rPD-1 exposure (Figure 1C). When cells were pre-treated for 30 minutes with an anti-PD-L1 blocking antibody LC3-II levels returned to baseline (Figure 1D; p<0.01). Results of experiments using mouse mammary 4T1 cells revealed similar LC3-II activation patterns, with LC3-II levels peaking at 30 minutes of incubation with rPD-1; this effect was also attenuated when cells were pre-treated with anti-PD-L1 Ab (Figure 1E).

**Figure 1:**
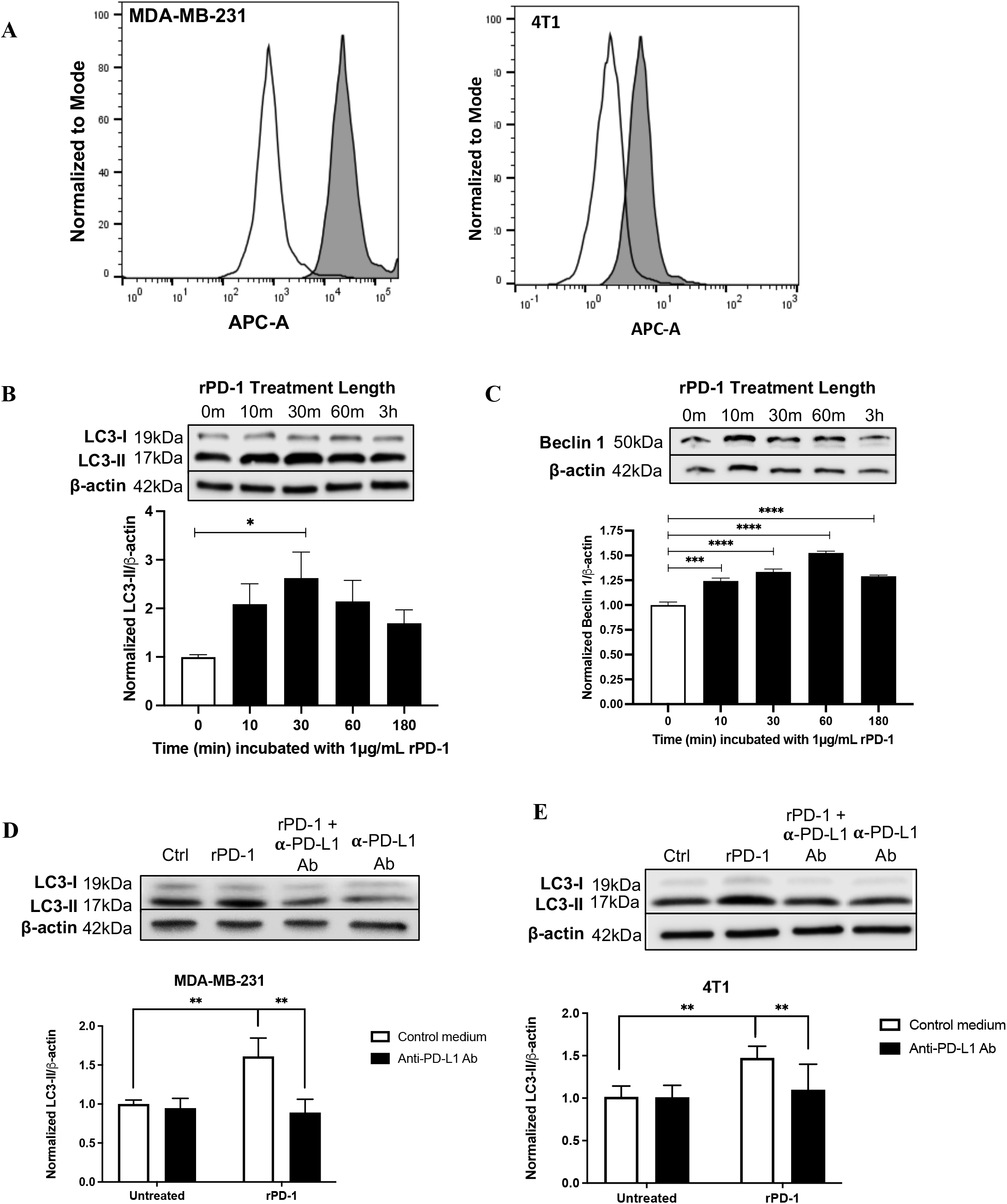
PD-1/PD-L1 reverse signaling increases expression of autophagy markers LC3-II and Beclin1. (A) PD-L1 expression of MDA-MB-231 and 4T1 cells was determined via flow cytometry. The shaded area represents staining with an anti-PD-L1 antibody, and the unshaded area represents staining with an isotype-matched control. MDA-MB-231 cells were incubated with 1 μg/mL rPD-1 and levels of (B) LC3-I, LC3-II and (C) Beclin1 protein were determined by western blotting. (D) MDA-MB-231 and (E) 4T1 cells were pre-treated with an anti-PD-L1 antibody and LC3-I to LC3-II conversion was assessed via western blot. The ratio of LC3-I and Beclin 1 to *β*-actin was quantified and compared between groups using a one-way ANOVA followed by Tukey’s multiple comparisons test. Results shown were pooled from at least three independent experiments *, *p*<0.05; ***,*p*<0.001.

### rPD-1 exposure increases the number of autophagosomes in human breast cancer cells

Upon activation of autophagy, lipidated LC3-II localizes to the autophagosome. To determine whether PD-1-induced PD-L1 signaling affects autophagosome formation, MDA-MB-231 cells were transfected with LC3 conjugated to GFP (GFP-LC3) and treated with rPD-1. Following 30 minutes of exposure to rPD-1, MDA-MB-231 cells displayed increased numbers of autophagosomes identified as GFP-positive punctate (Figure 2A). This increase in autophagosome numbers was comparable to the increase observed in cells cultured under serum starvation conditions, a stressor which induces autophagy. Furthermore, the increase in autophagosome numbers in rPD-1-treated cells was prevented by incubation with an anti-PD-L1 blocking antibody (Figure 2B).

**Figure 2:**
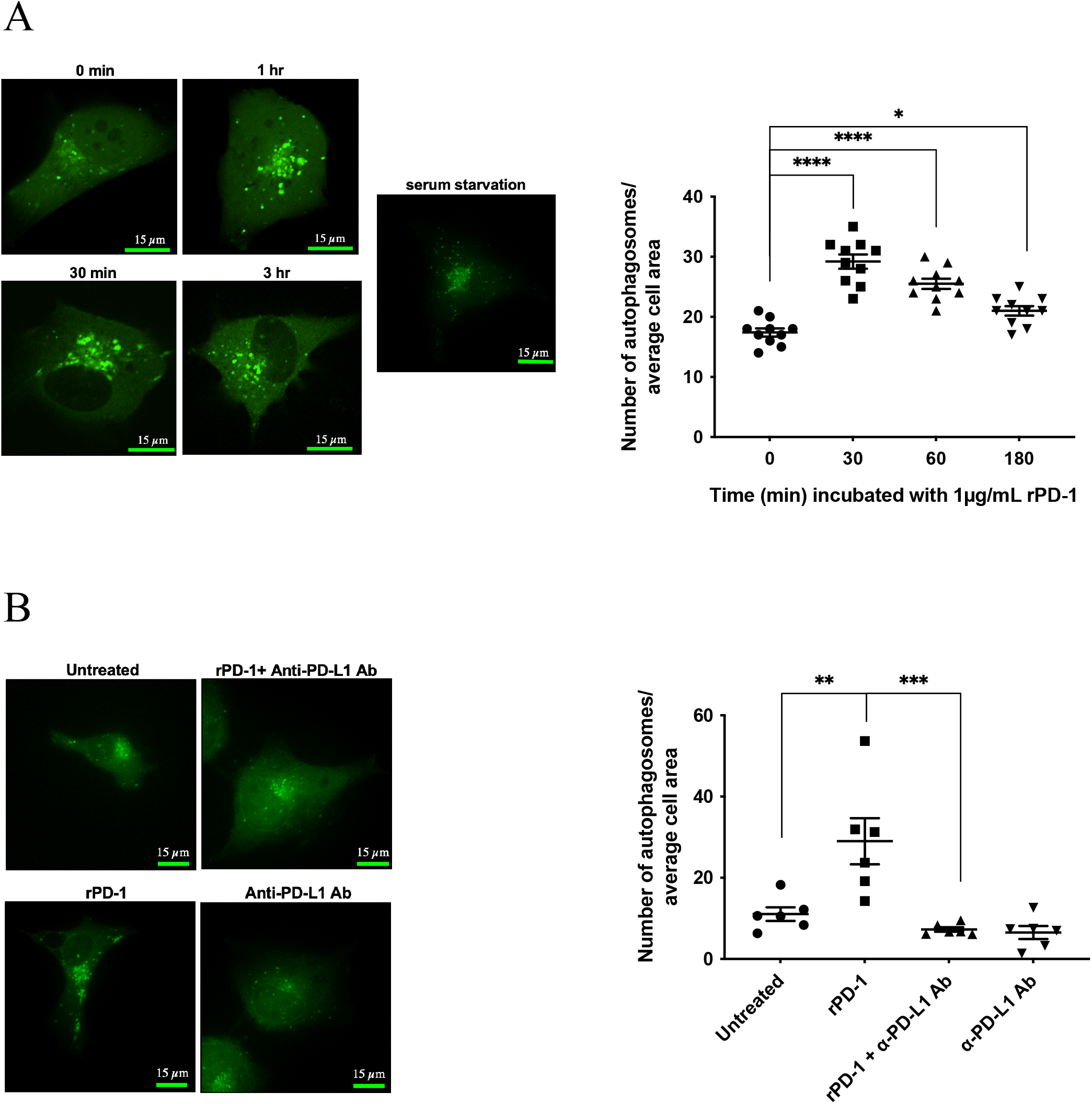
MDA-MB-231 cells display increased numbers of autophagosomes following treatment with rPD1. MDA-MB-231 cells transfected with GFP-LC3 were incubated with 1 μg/mL recombinant PD-1 for (A) 0, 30, 60, and 180 minutes or for (B) 30 min +/− an anti-PD-L1 antibody and autophagosomes were observed via immunofluorescence. Results were pooled from at least three independent experiments. *, *p*<0.05, **, *p*<0.01, ***, *p*<0.001.

### Inhibition of autophagy prevents PD-1/PD-L1-induced drug resistance

To assess the role of autophagy on PD-1/PD-L1 induced tumor cell drug resistance, we performed colony formation assays. Compared with control 4T1 cells incubated without rPD-1, incubation with rPD-1 did not affect the plating efficiency of these cells treated in the absence of doxorubicin, as shown by the lack of differences in their surviving fractions (Figure 3A-C). Incubation with doxorubicin decreased cell survival under all conditions (Figure 3A-C). However, pre-incubation with rPD-1 significantly increased the survival of control (scrambled shRNA) 4T1 cells following doxorubicin exposure (Figure 3A; p<0.01), but not the survival of Beclin 1- or Atg7-knockdown cells (Figure 3B and C).

**Figure 3:**
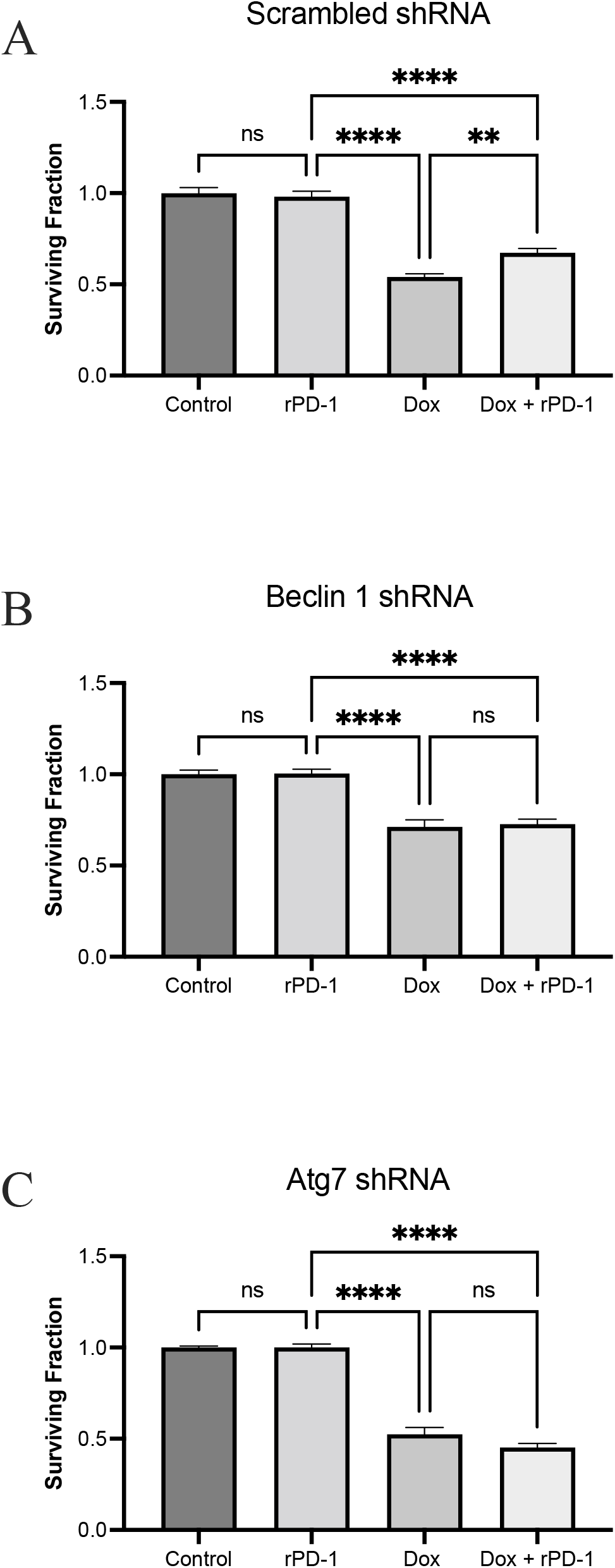
Knockdown of Beclin 1 and Atg7 abolishes PD-1-induced resistance to doxorubicin. Results of colony formation assays using 4T1 cells expressing scrambled shRNA (A), Beclin 1 shRNA (B), and Atg7 shRNA (C). Cells were incubated in the absence or presence of 1 μg/mL rPD-1 prior to exposure to 0.5 μM doxorubicin (or control medium) for one hour. Surviving fractions were calculated as described in *Materials and Methods.* Significance was determined by one-way ANOVA followed by Tukey’s multiple comparisons test. **, *p*<0.01, ****, *p*<0.0001.

### Autophagy mediated by PD-1/PD-L1 is dependent on ERK signaling

To elucidate potential upstream regulators of autophagy, we next examined whether activation of the PD-1/PD-L1 axis altered oncogenic signaling pathways. MDA-MB-231 cells treated with rPD-1 showed a time-dependent increase in phosphorylated ERK (pERK) (Figure 4A). To establish the role of ERK signaling in PD-1/PD-L1 induced autophagy, cells were treated with the MEK inhibitor U0126 (MEK is upstream of ERK and regulates its phosphorylation and activation) prior to rPD-1 treatment, as described. Immunoblot analysis of cells treated with U0126 confirmed inhibition of pERK phosphorylation (data not shown). Treatment with U0126 abolished the rPD-1 induced LC3-II accumulation in MDA-MB-231 cells (Figure 4B; p<0.01) as well as in 4T1 cells (Supplementary Figure 1; p<0.01). In addition, treatment of MDA-MB-231 cells with U0126 inhibited rPD-1 induced increase in autophagosome numbers (Figure 4C).

**Figure 4:**
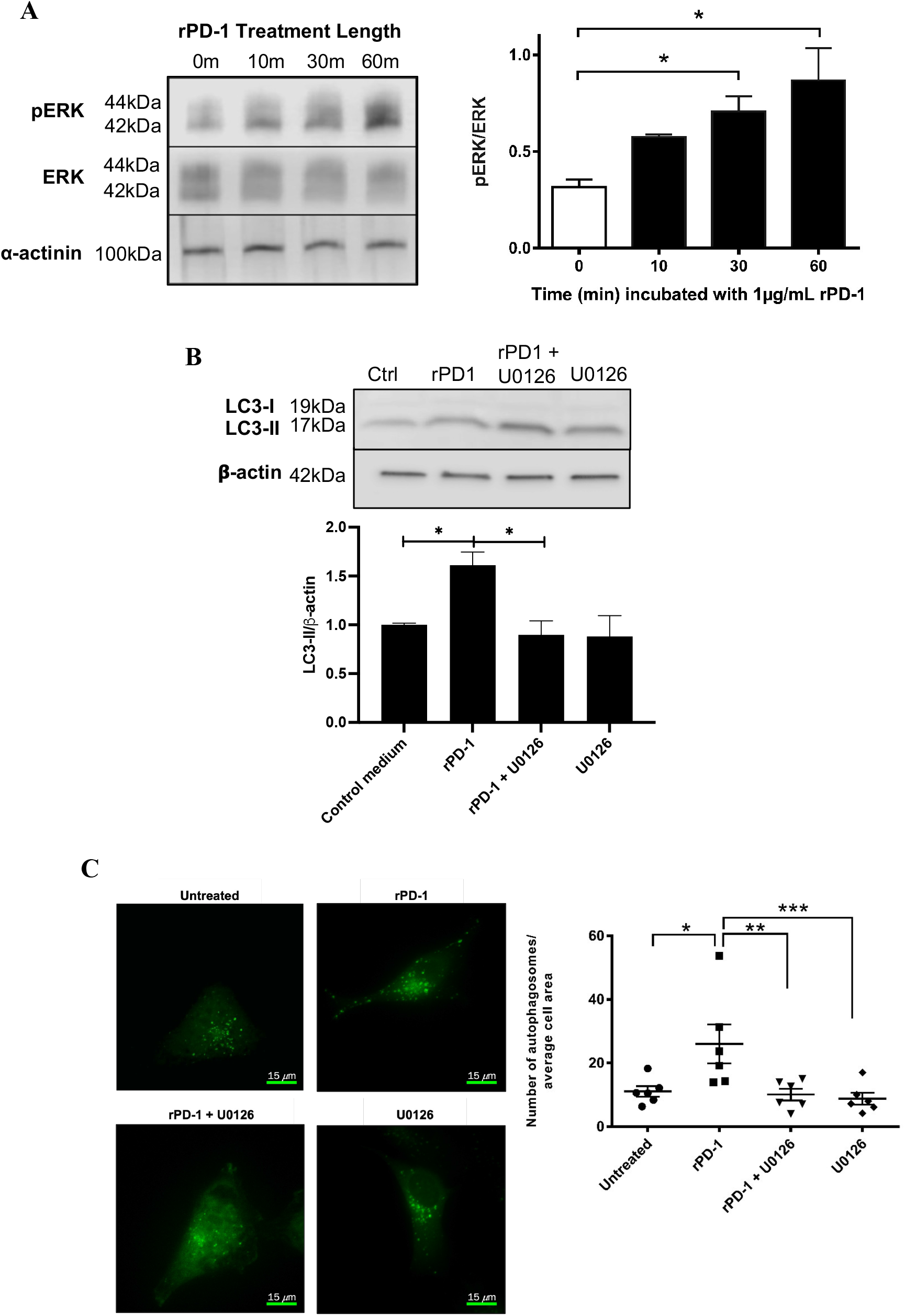
ERK inhibition with U0126 abolishes PD-1-induced autophagy in cancer cells. ERK inhibition with U0126 abolishes PD-1-induced autophagy in cancer cells. A. MDA-MB-213 cells were incubated with 1 μg/mL rPD-1 for 0, 10, 30, and 60 minutes and pERK expression was assessed via western blot. B. MDA-MB-213 were incubated with 1 μg/mL rPD-1 for 30 minutes ± 10 μM U0126 and LC3-I and LC3-II expression was assessed via western blot. C. MDA-MB-231 cells transfected with GFP-LC3 were incubated with 1 μg/mL recombinant PD-1 for 30 minutes ± 10 μM U0126 and autophagosome number was assessed via immunofluorescence. Significance was determined by one-way ANOVA followed byTukey’s post hoc test. Results shown were pooled from three independent experiments. *, P<0.05; **, P<0.01; ***, P<0.001. **.

## DISCUSSION

This study demonstrates a novel mechanism of drug resistance in tumor cells induced by reverse PD-1/PD-L1 signaling. Specifically, our results reveal that activation of PD-L1 signaling increases resistance to doxorubicin in mammary cancer cells via induction of autophagy. Furthermore, we provide evidence that oncogenic ERK signaling is a key mediator of autophagy induced by PD-1/PD-L1 signaling.

A study by Ishibashi *et al.* revealed that treating PD-L1-expressing myeloma cells with PD-1 stimulated drug resistance by inhibiting apoptotic pathways (7). Interestingly, Azuma *et al.* also demonstrated that tumor cells acquire resistance to pro-apoptotic mechanisms upon binding of PD-1 to PD-L1; however, these investigators provided evidence that classical apoptotic or anti-apoptotic pathways may not be involved in this process (6). Thus, when examining potential mechanisms of PD-1/PD-L1-mediated chemoresistance, we focused on the role of autophagy, as this biological process has been a proposed mediator of drug resistance (18–20). Our results indeed support a role for autophagy in PD-1/PD-L1-based chemoresistance. In contrast, a study by Clark *et al.* showed that low-PD-L1 expressing melanoma and ovarian cancer cells had decreased mammalian target of rapamycin (mTOR) activity, increased levels of autophagy, resistance to chloroquine (autophagy inhibitor) and slower tumor formation in immunocompromised mice compared to cells expressing normal levels of PD-L1 (21).

To further explore a causal role for autophagy in PD-1/PD-L1 induced drug resistance, we determined the effect of Beclin 1 and Atg7 knockdown on rPD-1 mediated resistance to doxorubicin. Beclin 1 is a component of class III PI3K complexes that produce phosphatidylinositol 3-phosphate [PI(3)P], which is essential for phagophore formation and the initiation of the autophagosome (20,22). Both Atg5 and Atg7 play important roles in the elongation and maturation of the autophagic membrane during canonical autophagy (23). However, cells can generate autophagic vacuoles via an alternative Atg5/Atg7-independent autophagy pathway (23). In our study, the fact that knockdown of either Beclin 1 or Atg7 resulted in the complete inhibition of rPD-1-mediated resistance to doxorubicin in 4T1 cells indicates that activation of the canonical autophagy pathway is required.

The Ras/Raf/MEK/ERK signal transduction pathway is an important mediator of malignant phenotypes and ERK activation has been shown to lead to both apoptotic and autophagic cell death in various models (24–26). In contrast, inhibition of ERK phosphorylation has been associated with decreased autophagy and increased susceptibility to tumor necrosis factor (TNF)-induced cell death (27). Furthermore, ERK activation induces conversion of LC3-I to LC3-II, a key step in autophagosome formation (28). Our study established that PD-1/PD-L1 reverse signaling leads to increased phosphorylation of ERK, implicating this pathway as a potential driver of PD-1/PD-L1 induced autophagy and drug resistance. Furthermore, blocking this signaling pathway with a MEK inhibitor, U0126, led to a decrease in rPD-1-induced LC3-II expression and autophagosome numbers. Interestingly, it has been shown that activation of Raf, an upstream mediator of ERK signaling, leads to resistance to doxorubicin and paclitaxel in breast cancer cells (29). The relative contribution of autophagy to Raf-mediated drug resistance warrants further investigation. The phosphoinositide 3-kinase (PI3K)/protein kinase B (AKT) signaling pathway has also been implicated in PD-1/PD-L1 induced drug resistance; however, this pathway may only be involved in an antiapoptotic response (7). The PI3K/AKT signaling pathway is also known to inhibit autophagy (30,31), thus it is possible that it plays a role in regulating PD-1/PD-L1 induced drug resistance.

In addition to mediating chemoresistance, autophagy has implications for tumor progression as it is a mechanism of stress response. It has been shown in multiple cancers that high levels of autophagy are required for the maintenance of metabolic function, and that inhibition of autophagy increases the intracellular concentrations of reactive oxygen species and leads to DNA damage (32,33). Therefore, PD-1/PD-L1-mediated induction of autophagy in tumor cells may promote tumor progression and malignancy in addition to inducing drug resistance. Furthermore, ERK signaling is important for the acquisition of malignant phenotypes, including cell proliferation, migration, and differentiation. This suggests that PD-1/PD-L1 reverse signaling may not only promote drug resistance, but also the acquisition of other malignant phenotypes.

## CONCLUSIONS

The widespread expression of PD-L1, its role as a prognostic marker for cancer outcomes, and the unprecedented clinical effects of anti-PD-1/PD-L1 treatment, make the characterization of this axis critical to improving on current therapeutics. This study presents autophagy via ERK signaling as a potential mechanism for PD-1/PD-L1 induced drug resistance. Chemoresistance is still an important barrier to cancer treatment and elucidating mechanisms of drug resistance is essential for improving therapeutic efficacy. Our study suggests that there is a potential role for combining autophagy inhibitors, as well as Ras/Raf/MEK/ERK inhibitors with PD-1/PD-L1 blocking agents and standard chemotherapy protocols. Chloroquine, an inhibitor of autophagy, has been used in combination therapies resulting in enhanced efficacy of tumor cell killing. There are also FDA approved inhibitors of Raf (dabrafenib) and MEK (trametinib), as well as various ERK inhibitors currently in clinical trials. Combining these therapies may sensitize tumor cells to current gold standard treatments and improve patient outcomes.

## DECLARATIONS

### Competing Interests

None of the authors declare any potential conflict of interest.

### Funding

This work was supported by a project grant from the Canadian Institutes of Health Research (PJT-148601) awarded to C.H. Graham, M. Koti, and D.R Siemens. L. Minassian was the recipient of a Canadian Institutes of Health Research Doctoral Award-Frederick Banting and Charles Best Canada Graduate Scholarship. D. Sanwalka received funding from the Terry Fox Research Institute Transdisciplinary Training Program in Cancer Research and T. Cotechini was a recipient of a Canadian Institutes of Health Research Post-Doctoral Fellowship.

### Authors’ Contributions

LM and DS were involved in study conceptualization and design, performed all experiments and data analysis, and wrote the original manuscript. SMG, LS, JFP, CLM and AG aided in experimental procedures. DRS, MK, EC, and AWBC provided conceptual advice, as well as technical and material support. TC edited the manuscript. CHG was involved in study conceptualization and analysis of data, supervised the execution of the study, edited the manuscript, and acquired funding to perform this research. All authors read and approved the manuscript

## Acknowledgments

We would like to thank Brooke Ring for her assistance with flow cytometry experiments. We would also like to thank Matt Gordon for aiding with the autophagosome fluorescence imaging.

## LIST OF ABBREVIATIONS

Ab: antibody
AKT: protein kinase B
ANOVA: analysis of variance
APC: allophycocyanin
ATCC: American Type Culture Collection
Atg7: autophagy-related gene 7
ERK: extracellular signal-regulated kinase
FBS: fetal bovine serum
GFP: green fluorescence protein
MEK: mitogen-activated protein kinase kinase
mTOR: mammalian target of rapamycin
PD-1: programmed cell death protein-1
PD-L1: programmed death ligand-1
PE: plating efficiency
PI3K: phosphoinositide 3-kinase
PVDF: polyvinylidene difluoride
rPD-1: recombinant PD-1
RPMI: Roswell Park Memorial Institute
SDS-PAGE: sodium dodecyl sulphate-polyacrylamide gel electrophoresis
SF: surviving fraction
TBST: tris-buffered saline, 0.1% Tween 20
TNF: tumor necrosis factor

## FIGURE LEGENDS

**Supplementary Figure 1:**
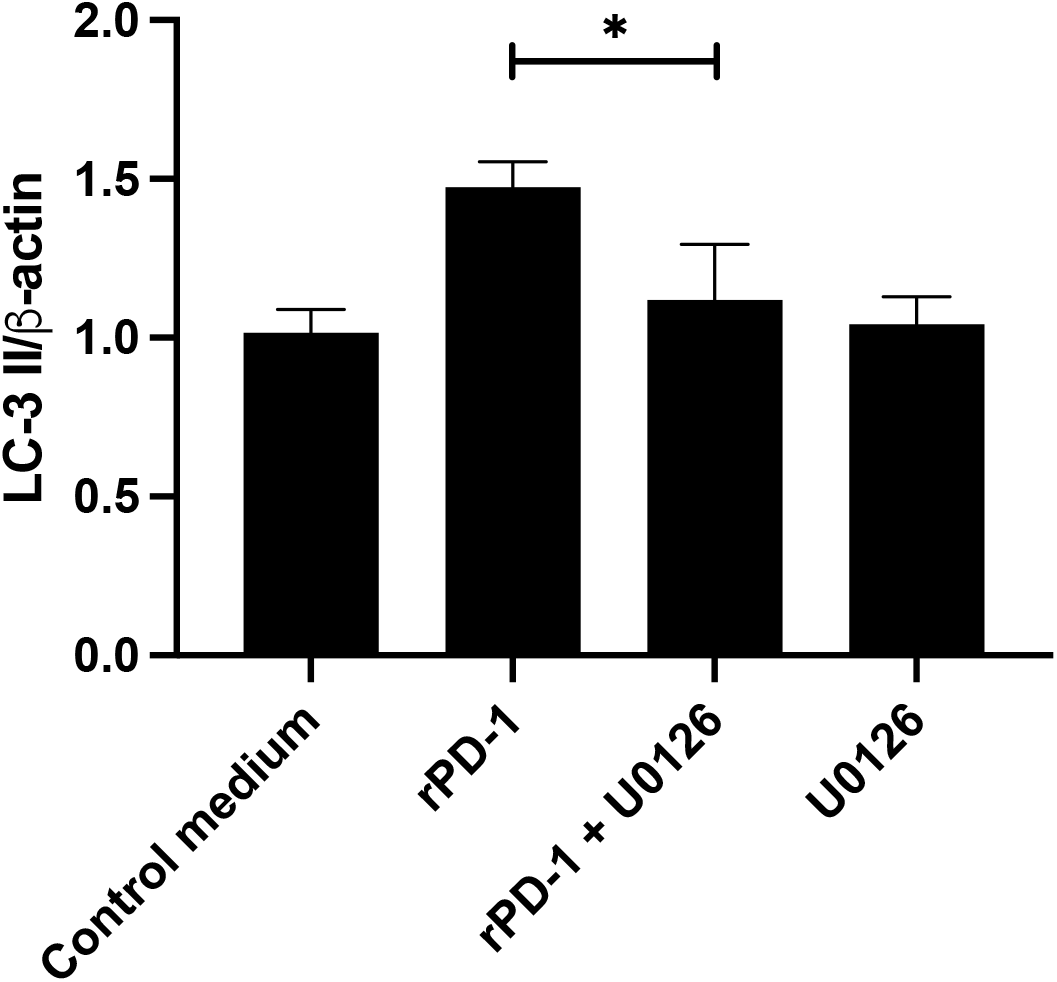
U0126 inhibits rPD-1 mediated LC-3II accumulation in 4T1 cells. 4T1 were incubated with 1 μg/mL rPD-1 for 30 minutes ± 10 μM U0126 and LC3-I and LC3-II expression was assessed by western blot. *, p<0.05.

## Notes

### Competing Interest Statement

The authors have declared no competing interest.

## REFERENCES

1. Chen L, Han X. Anti-PD-1/PD-L1 therapy of human cancer: past, present, and future. J Clin Invest 2015;125:3384–91

2. Swaika A, Hammond WA, Joseph RW. Current state of anti-PD-L1 and anti-PD-1 agents in cancer therapy. Mol Immunol 2015;67:4–17

3. Gong J, Chehrazi-Raffle A, Reddi S, Salgia R. Development of PD-1 and PD-L1 inhibitors as a form of cancer immunotherapy: a comprehensive review of registration trials and future considerations. J Immunother Cancer 2018;6:8

4. Ferrara R, Pilotto S, Caccese M, Grizzi G, Sperduti I, Giannarelli D, et al. Do immune checkpoint inhibitors need new studies methodology? J Thorac Dis 2018;10:S1564–S80

5. Kubli SP, Berger T, Araujo DV, Siu LL, Mak TW. Beyond immune checkpoint blockade: emerging immunological strategies. Nat Rev Drug Discov 2021;20:899–919

6. Azuma T, Yao S, Zhu G, Flies AS, Flies SJ, Chen L. B7-H1 is a ubiquitous antiapoptotic receptor on cancer cells. Blood 2008;111:3635–43

7. Ishibashi M, Tamura H, Sunakawa M, Kondo-Onodera A, Okuyama N, Hamada Y, et al. Myeloma Drug Resistance Induced by Binding of Myeloma B7-H1 (PD-L1) to PD-1. Cancer Immunol Res 2016;4:779–88

8. Black M, Barsoum IB, Truesdell P, Cotechini T, Macdonald-Goodfellow SK, Petroff M, et al. Activation of the PD-1/PD-L1 immune checkpoint confers tumor cell chemoresistance associated with increased metastasis. Oncotarget 2016;7:10557–67

9. Li X, Zhou Y, Li Y, Yang L, Ma Y, Peng X, et al. Autophagy: A novel mechanism of chemoresistance in cancers. Biomed Pharmacother 2019;119:109415

10. Mathew R, Kongara S, Beaudoin B, Karp CM, Bray K, Degenhardt K, et al. Autophagy suppresses tumor progression by limiting chromosomal instability. Genes Dev 2007;21:1367–81

11. Kondo Y, Kanzawa T, Sawaya R, Kondo S. The role of autophagy in cancer development and response to therapy. Nat Rev Cancer 2005;5:726–34

12. Liu D, Yang Y, Liu Q, Wang J. Inhibition of autophagy by 3-MA potentiates cisplatin-induced apoptosis in esophageal squamous cell carcinoma cells. Med Oncol 2011;28:105–11

13. Zou Z, Yuan Z, Zhang Q, Long Z, Chen J, Tang Z, et al. Aurora kinase A inhibition-induced autophagy triggers drug resistance in breast cancer cells. Autophagy 2012;8:1798–810

14. Sui X, Chen R, Wang Z, Huang Z, Kong N, Zhang M, et al. Autophagy and chemotherapy resistance: a promising therapeutic target for cancer treatment. Cell Death Dis 2013;4:e838

15. Sun WL, Chen J, Wang YP, Zheng H. Autophagy protects breast cancer cells from epirubicin-induced apoptosis and facilitates epirubicin-resistance development. Autophagy 2011;7:1035–44

16. Nwadike C, Williamson LE, Gallagher LE, Guan JL, Chan EYW. AMPK Inhibits ULK1-Dependent Autophagosome Formation and Lysosomal Acidification via Distinct Mechanisms. Mol Cell Biol 2018;38

17. Gallagher LE, Radhi OA, Abdullah MO, McCluskey AG, Boyd M, Chan EYW. Lysosomotropism depends on glucose: a chloroquine resistance mechanism. Cell Death Dis 2017;8:e3014

18. Honscheid P, Datta K, Muders MH. Autophagy: detection, regulation and its role in cancer and therapy response. Int J Radiat Biol 2014;90:628–35

19. Yang ZJ, Chee CE, Huang S, Sinicrope FA. The role of autophagy in cancer: therapeutic implications. Mol Cancer Ther 2011;10:1533–41

20. Zamame Ramirez JA, Romagnoli GG, Kaneno R. Inhibiting autophagy to prevent drug resistance and improve anti-tumor therapy. Life Sci 2021;265:118745

21. Clark CA, Gupta HB, Sareddy G, Pandeswara S, Lao S, Yuan B, et al. Tumor-Intrinsic PD-L1 Signals Regulate Cell Growth, Pathogenesis, and Autophagy in Ovarian Cancer and Melanoma. Cancer Res 2016;76:6964–74

22. He C, Levine B. The Beclin 1 interactome. Curr Opin Cell Biol 2010;22:140–9

23. Urbanska K, Orzechowski A. The Secrets of Alternative Autophagy. Cells 2021;10

24. Cagnol S, Chambard JC. ERK and cell death: mechanisms of ERK-induced cell death--apoptosis, autophagy and senescence. FEBS J 2010;277:2–21

25. Sun Y, Liu WZ, Liu T, Feng X, Yang N, Zhou HF. Signaling pathway of MAPK/ERK in cell proliferation, differentiation, migration, senescence and apoptosis. J Recept Signal Transduct Res 2015;35:600–4

26. Woessmann W, Chen X, Borkhardt A. Ras-mediated activation of ERK by cisplatin induces cell death independently of p53 in osteosarcoma and neuroblastoma cell lines. Cancer Chemother Pharmacol 2002;50:397–404

27. Sivaprasad U, Basu A. Inhibition of ERK attenuates autophagy and potentiates tumour necrosis factor-alpha-induced cell death in MCF-7 cells. J Cell Mol Med 2008;12:1265–71

28. Oh SH, Lim SC. Endoplasmic reticulum stress-mediated autophagy/apoptosis induced by capsaicin (8-methyl-N-vanillyl-6-nonenamide) and dihydrocapsaicin is regulated by the extent of c-Jun NH2-terminal kinase/extracellular signal-regulated kinase activation in WI38 lung epithelial fibroblast cells. J Pharmacol Exp Ther 2009;329:112–22

29. McCubrey JA, Steelman LS, Chappell WH, Abrams SL, Wong EW, Chang F, et al. Roles of the Raf/MEK/ERK pathway in cell growth, malignant transformation and drug resistance. Biochim Biophys Acta 2007;1773:1263–84

30. Wang RC, Wei Y, An Z, Zou Z, Xiao G, Bhagat G, et al. Akt-mediated regulation of autophagy and tumorigenesis through Beclin 1 phosphorylation. Science 2012;338:956–9

31. Degtyarev M, De Maziere A, Orr C, Lin J, Lee BB, Tien JY, et al. Akt inhibition promotes autophagy and sensitizes PTEN-null tumors to lysosomotropic agents. J Cell Biol 2008;183:101–16

32. Yang S, Wang X, Contino G, Liesa M, Sahin E, Ying H, et al. Pancreatic cancers require autophagy for tumor growth. Genes Dev 2011;25:717–29

33. Guo JY, Chen HY, Mathew R, Fan J, Strohecker AM, Karsli-Uzunbas G, et al. Activated Ras requires autophagy to maintain oxidative metabolism and tumorigenesis. Genes Dev 2011;25:460–70

